# Acetate as a metabolic booster for glucose-based bioproduction in Escherichia coli

**DOI:** 10.1101/2025.06.17.659982

**Authors:** Thomas Gosselin-Monplaisir, Denis Jallet, Erwana Harscoet, Sandrine Uttenweiler-Joseph, Stéphanie Heux, Pierre Millard

## Abstract

*Escherichia coli*, a key workhorse in industrial biotechnology, is commonly grown on glucose, which supports rapid growth but leads to acetate overflow, which instead inhibits growth, diverts carbon from the production pathway, and reduces productivity. Recent studies suggest that acetate may also have beneficial effects in glucose-grown *E. coli*, but its potential in bioprocesses remains unexplored. In this study, we systematically investigated acetate’s impact on bioproduction using a kinetic model of glucose and acetate metabolism in *E. coli*. The model predicts that acetate can enhance bioproduction in glucose-grown *E. coli* through three mechanisms: (i) by minimizing acetate overflow, thereby reducing carbon loss, (ii) by increasing acetyl-CoA levels, thereby boosting the biosynthetic flux of acetyl-CoA-derived compounds, and (iii) by promoting biomass accumulation, thus improving overall productivity. We experimentally validated the predictions of the model for mevalonate and 3-hydroxypropionate production, where acetate supplementation increased productivity by 117% and 34%, respectively. Our findings provide a valuable framework for optimizing *E. coli*-based bioprocesses and highlight acetate’s underutilized potential in biotechnology. By leveraging acetate from waste streams as a metabolic booster, this approach could contribute to more sustainable and environmentally friendly bioprocesses.

**Highlights:**

- A kinetic model was used for rational optimization of *E. coli*-based bioprocesses
- Acetate can enhance growth and production of acetyl-CoA-derived bioproducts
- Model predictions were validated for mevalonate and 3-hydroxypropionate production
- Acetate can be used as a metabolic booster for glucose-based bioproduction in *E. coli*

## 1. Introduction

The genetic tractability, rapid growth, and metabolic versatility of *Escherichia coli* have established it as the primary chassis organism in biotechnology. *E. coli* plays a pivotal role in industrial biotechnology, supporting applications in biofuels, pharmaceuticals, bioplastics, waste recycling, and enzyme production (Kim et al., 2025; Pontrelli et al., 2018; Vickers et al., 2012). Advances in synthetic biology and metabolic engineering continue to enhance its capabilities, making it one of the most important microbial platforms for industrial applications.

The primary carbon sources in *E. coli*-based bioprocesses are glycolytic substrates such as glucose and glycerol, chosen for their availability, metabolic efficiency, and cost-effectiveness (Martínez-Gómez et al., 2012; Pontrelli et al., 2018; Wang et al., 2015). However, a major limitation of fast glycolytic growth in *E. coli* is the co-production of undesirable metabolites, primarily acetate, through overflow metabolism (De Mey et al., 2007). Acetate is traditionally considered detrimental to bioprocess efficiency because of its inhibitory effects on growth and because it diverts carbon away from bioproduction pathways (De Mey et al., 2007; Eiteman and Altman, 2006).

Current strategies to optimize *E. coli*-based bioprocesses focus on minimizing acetate accumulation by strain engineering or bioprocess control (De Mey et al., 2007; Gecse et al., 2024). Strain engineering can be used to (i) modify acetate metabolism to reduce acetate formation or enable acetate reuptake (Phue et al., 2010; Tao et al., 2012; H.-D. Wang et al., 2023), (ii) redirect metabolic flux away from the acetate pathway and into the TCA cycle (Waegeman et al., 2011; L. Wang et al., 2023; Zhu et al., 2019), and/or (iii) reduce glucose uptake to prevent acetate overflow (Fuentes et al., 2013; Zhu et al., 2019). Acetate accumulation can also be mitigated by limiting glucose uptake through fermentation strategies, such as chemostat or fed-batch cultures (De Mey et al., 2007; Gecse et al., 2024; Kang et al., 2024). However, these strategies may alter cellular metabolism and are often counterproductive (De Mey et al., 2007). For instance, overexpressing acetyl-CoA synthetase to enable acetate reuptake creates a futile cycle that wastes ATP, making it energetically costly for the cell (Valgepea et al., 2010).

Despite its negative effect on glycolytic-based bioprocess efficiency, acetate can also serve as a primary carbon source for *E. coli*, albeit with slow growth and low productivity when it is the only carbon source (Chong et al., 2013; Gong et al., 2022; Noh et al., 2018). Nevertheless, acetate is a promising feedstock, metabolic intermediate, and industrial precursor for biofuels, bioplastics, pharmaceuticals, and synthetic biology applications (Kiefer et al., 2021). Acetate is particularly interesting for the production of acetyl-CoA-derived compounds (Gong et al., 2022; Kiefer et al., 2021; Lama et al., 2021; Zhang et al., 2022), as acetyl-CoA is a highly versatile precursor at the intersection of glycolytic and acetate metabolisms. Since acetate can be sourced from CO_2_ (H. Wang et al., 2023) and various waste streams, including agricultural and industrial by-products, its potential role in carbon recycling and sustainable bioprocessing is also attracting increasing attention (Kiefer et al., 2021; Zhang et al., 2022).

Another interesting property of acetate in *E. coli* is that its negative and positive effects are not mutually exclusive. Recent studies have shown that acetate can be co-utilized with glycolytic substrates (Enjalbert et al., 2017; Millard et al., 2023, 2021), potentially accelerating bioprocesses by enhancing the production of acetyl-CoA-derived compounds. While no study has yet investigated this mechanism in detail, co-utilization of acetate with glucose has been shown to improve butyl-butyrate production by *E. coli* (Zhang et al., 2024), hyaluronic acid production by *Lactococcus lactis* (Puvendran and Jayaraman, 2019), and itaconic acid production by *Ustilago maydis* (Ziegler et al., 2024). Acetate overflow can also provide a competitive advantage to *E. coli* under certain conditions (Rabbers et al., 2022) and serve as a metabolic valve, helping maintain metabolic balance and flexibility to environmental changes (Enjalbert et al., 2017; Valgepea et al., 2010). Furthermore, recent findings indicate that acetate is not always toxic to *E. coli* during growth on glycolytic substrates (Millard et al., 2023). At low glycolytic flux, acetate may increase metabolic robustness to glycolytic perturbations and even enhance growth (Millard et al., 2023), highlighting an underexplored opportunity to harness its full potential in biotechnology.

In this study, we systematically investigated the potential of acetate in *E. coli*-based bioproduction of acetyl-CoA-derived compounds from glucose. We first used kinetic modeling to evaluate *E. coli*’s response to acetate *in silico* and then experimentally validated the model’s predictions, using mevalonate and 3-hydroxypropionate (3HP), two important chemicals in industrial biotechnology, as test cases.

## 2. Materials and Methods

### 2.1. Strain construction

The strains and plasmids used in this study are listed in Table 1. The parent strain was *E. coli* K-12 BW25113.

**Table 1.** Strains and plasmids used in this study.

| Strain or plasmid | Genotype or description | Source |
| --- | --- | --- |
| **Plasmids** |  |  |
| pMevT | pMevT, Addgene plasmid # 17815 ;<br>http://n2t.net/addgene:17815 ; RRID:Addgene 17815.<br>ChlR | (Martin et al., 2003) |
| pMevE | pMevT has been digested by HindIII and ligated to<br>generate the empty plasmid. ChlR | This study |
| p3HP | pSEVA131 with *mcr* gene from *Chloroflexus aurantiacus* (GenBank: AAS20429, codon-optimized) under the control of an IPTG-inducible promoter (Ptrc) has been synthesized and subcloned by GeneCust. AmpR | This study |
| pSEVA131 | Detailed in the reference. AmpR | (Martínez-García et al., 2023) |
| **Bacterial strains** |  |  |
| WT | *E. coli* K-12 BW25113 WT [F-λ- ΔlacZ4787(::rrnB-3) hsdR514 $\Delta$(araBAD)567 $\Delta$(rhaBAD)568] | Keio collection (Baba et al., 2006) |
| Δ*pgi* | *E. coli* K-12 BW25113 Δ*pgi* strain from Keio collection (Baba et al., 2006) cured of kanamycin cassette (Datsenko and Wanner, 2000) | This study |

### 2.2. Cultures

*E. coli* was grown in M9 minimal media (Millard et al., 2014) supplemented with 15 mM glucose. Sodium acetate (prepared as a 1.2 M solution at pH 7.0 by addition of potassium hydroxide) was added up to the required concentration, and IPTG (prepared as a 250 mM solution) was added at a final concentration of 1 mM. For isotope labeling experiments, unlabeled acetate was replaced by U-^13^C_2_-acetate (Eurisotop, France). The cells were grown in 250 mL shake flasks at 37°C and 200 rpm, in 20 mL of medium. Growth was monitored by measuring the optical density (OD) at 600 nm using a Genesys 150 ThermoScientific spectrophotometer. Biomass concentration was determined using a conversion factor of 0.42 g_CDW_/L/OD unit.

### 2.3. Metabolomics

#### Exometabolome analysis

Extracellular concentrations of glucose, acetate, mevalonate, and 3-hydroxypropionate were quantified during growth by 1D ^1^H-NMR on a Bruker 500 MHz spectrometer equipped with a 5 mm TXI cryoprobe (Bruker, Germany). Samples (180 µL) of filtered broth (0.2 μm, Sartorius, Germany) were mixed with 20 µL of D_2_O containing 10 mM TSPd4 (deuterated 3-(trimethylsilyl)-1-propanesulfonic acid, used as internal standard). The ZGPR presaturation sequence was used for water signal suppression, with a 30° pulse and a 10 s relaxation delay between scans to ensure full signal recovery. A total of 32 scans were accumulated (64 k data points with a spectral width of 16 ppm) after 4 dummy scans. Spectra were processed using TopSpin v4.1.4 (Bruker).

#### Intracellular acetyl-CoA quantification

Cells were separated from the medium by fast filtration. Filters were washed with 1 mL of fresh medium containing the same carbon sources as the growth medium, quickly transferred into liquid nitrogen, and stored at −80 °C until metabolite extraction. Intracellular metabolites were extracted by incubating the filters for 1 h in 5 mL of extraction solution (MeOH:ACN:H_2_O solution 40:40:20 with 0.5% formic acid) at −20°C, after adding 50 µL of a fully ^13^C-labeled internal standard. The extracts were centrifuged at 12,000 g for 5 min, and the supernatants were evaporated under vacuum in a SpeedVac (Savant SC250EXP) overnight. The dried extracts were then resuspended in 200 µL of milliQ water. Samples were analyzed by liquid chromatography on a Dionex ICS 5000 system coupled to a high-resolution Q-Exactive Plus mass spectrometer, as detailed in (Lajus et al., 2020). The raw data were processed using Skyline v24.1.0.

#### Isotopic analyses of mevalonate

The ^13^C-enrichment of extracellular mevalonate was measured in exometabolome samples (diluted 1000 times) using ion chromatography on a Dionex ICS-5000 coupled to a high-resolution Q-Exactive Plus mass spectrometer, as detailed in (Pineda et al., 2018). The intensities of each mass fraction in the isotopic cluster of mevalonate were extracted from the raw data using Skyline v24.1.0. The mean ^13^C-enrichment was calculated from these mass fractions after correction for naturally occurring isotopes and the isotopic purity of the ^13^C-labeled acetate (99%), using IsoCor v2.2.2 (Millard et al., 2019) (https://github.com/MetaSys-LISBP/IsoCor).

### 2.4. Extracellular flux calculations

Extracellular fluxes (i.e. glucose uptake, acetate flux, mevalonate production, 3HP production, and growth rates) were determined from the concentration time courses of biomass, substrates, and products using PhysioFit v3.4.0 (Le Grégam et al., 2024) (https://github.com/MetaSys-LISBP/PhysioFit).

### 2.5. Kinetic modeling

We used a published kinetic model of *E. coli* (Millard et al., 2021), available from the Biomodels database (https://www.ebi.ac.uk/biomodels; (Chelliah et al., 2015)) under identifier MODEL2005050001. This model was extended to include a bioproduction pathway converting acetyl-CoA into an extracellular product, with irreversible Michaelis-Menten kinetics (with V_max_ set to 10% of the maximal glucose uptake rate, and K_M_ = 5 mM or 5 µM). The models used in this study are available from the Biomodels database under identifiers MODEL2506160001 and MODEL2506160002. Simulations were performed as described in the text, using COPASI (Hoops et al., 2006), v4.39) with the COPASI R Connector (CoRC) package (v0.12, https://github.com/jpahle/CoRC, (Förster et al., 2021)) in R (v4.4.2, https://www.r-project.org). The R notebook used to run the simulations and generate the figures is provided at https://github.com/MetaSys-LISBP/glucose_acetate_bioproduction.

### 2.6. Statistical analyses

Results were compared between strains or conditions using two-sided t-tests.

## 3. Results

### 3.1. Model-guided strategy to evaluate acetate’s effect on bioproduction

We used kinetic modeling to systematically explore whether acetate can enhance *E. coli* glucose-based bioproduction. We incorporated a production pathway converting acetyl-CoA into a product, P, into an extensively validated model of glucose and acetate metabolism (Millard et al., 2021, 2023; Figure 1).

**Figure 1.**
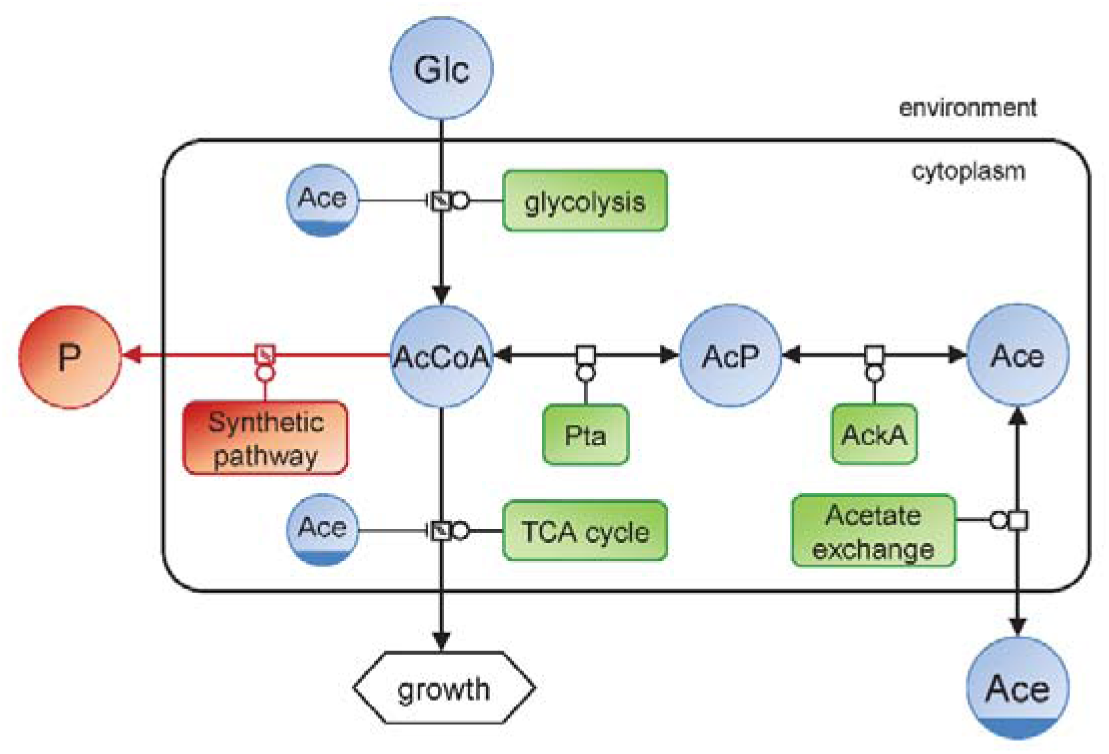
Representation (in Systems Biology Graphical Notation format, http://sbgn.org) of the glucose and acetate metabolism of an *E. coli* strain engineered to produce a product P, as implemented in the kinetic model used in the study. The native metabolism is shown in blue (metabolites) and green (enzymes), and the synthetic bioproduction pathway is shown in red.

Enzyme saturation is a key determinant of metabolic efficiency, shaping flux control, resource allocation, and pathway optimality (Noor and Liebermeister, 2024). To capture this essential feature, we modeled the production reaction as a Michaelis-Menten rate law, with V_max_ arbitrarily set to 10% of the maximal glucose uptake rate to represent a bioproduction pathway with limited enzymatic capacity. Enzyme saturation, defined by the Michaelis constant (K_M_), dictates how metabolic flux responds to intracellular metabolite levels (Castaño-Cerezo et al., 2024). At low substrate concentrations, the flux is responsive to substrate fluctuations, allowing for dynamic regulation but at the cost of inefficient enzyme utilization. Conversely, at high substrate concentrations, the pathway operates at maximum capacity, ensuring stable fluxes and optimal enzyme utilization, but with reduced flexibility and potential metabolic imbalances. This trade-off is critical in bioproduction, where balancing pathway flexibility and robustness is key to optimizing process efficiency.

To capture these differences, we modeled two configurations of the production pathway with different levels of enzyme saturation by adjusting the K_M_ of the production pathway. Given that the intracellular acetyl-CoA concentration in glucose-grown *E. coli* is typically between 0.02 and 0.6 mM (Bennett et al., 2009; Chohnan et al., 1998; Takamura et al., 1985), a high K_M_ (5 mM) was used to simulate a pathway operating below substrate saturation, making it sensitive to changes in acetyl-CoA concentration, while a low K_M_ (5 µM) was used to model the near-saturation state, with flux largely independent of acetyl-CoA fluctuations. Tuning pathway saturation allowed us to strategically explore the trade-off between pathway flexibility and robustness and to determine the impact of acetate on the balance between production and growth.

We simulated *E. coli* growth on glucose (15 mM) in both configurations for a wide range of acetate concentrations (0.1 to 100 mM) and glycolytic flux levels (20% to 100% of the maximal glycolytic flux). This allowed us to predict how acetate influences cell physiology (growth rate, acetate and glucose fluxes, intracellular acetyl-CoA levels) and bioprocess performance (product flux, yield, and productivity – three key metrics of bioprocess success). In the following sections, we present the results of the model for both configurations, along with those of a test case designed to evaluate the model’s predictions.

### 3.2. Predictions for an acetyl-CoA-sensitive production pathway

With the acetyl-CoA-sensitive production pathway (K_M_ = 5 mM), the predicted trends in glucose uptake rates (Figure 2A), acetate fluxes (Figure 2B), and growth rates (Figure 2C) under different environmental conditions are similar to those observed in the non-producing *E. coli* strain, namely, (i) glucose uptake decreases at increasing acetate concentrations, (ii) the acetate flux reverses above a certain acetate concentration, a threshold that depends on the glycolytic flux, and (iii) growth rates are enhanced by acetate at low glucose uptake rates. Additional details on these predictions, which have been experimentally validated in a non-producing *E. coli* strain, can be found in (Millard et al., 2023).

**Figure 2.**
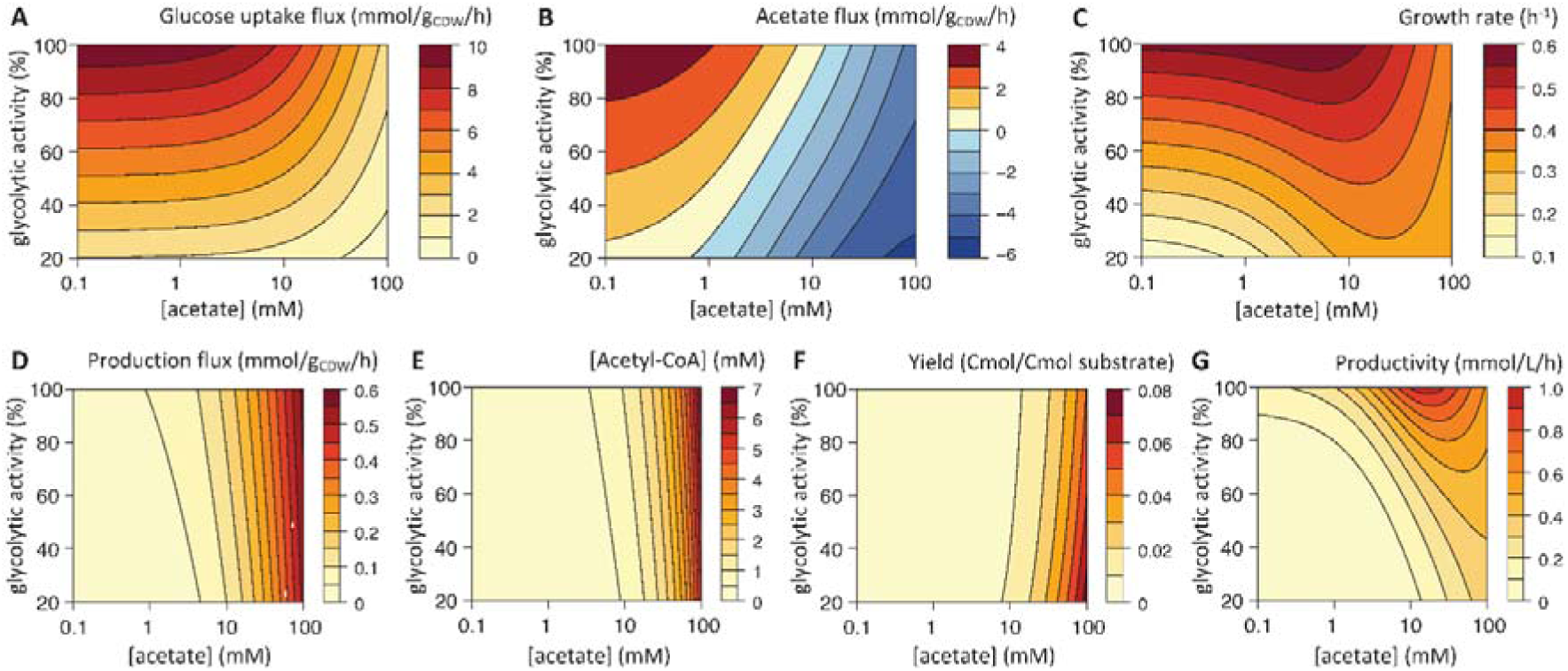
Predicted response to glycolytic and acetate perturbations of an *E. coli* strain with an acetyl-CoA-sensitive production pathway. Glucose uptake flux (A), acetate flux (B), growth rate (C), product flux (D), intracellular acetyl-CoA concentration (E), product yield (F) and volumetric productivity (G) simulated over a broad range of glycolytic activity levels (20 to 100% of the initial V_max_) and acetate concentrations (0.1 to 100 mM).

The model predicts that increasing the acetate concentration should enhance the production flux (Figure 2D) due to an increase in the intracellular concentration of acetyl-CoA (Figure 2E). However, the enzyme is not used to full capacity: the production flux varies considerably but only reaches at most 60% of the maximal rate (0 to 0.6 mmol/g_CDW_/h compared with V_max_ = 1 mmol/g_CDW_/h). According to the model, acetate should improve the product yield (Figure 2F), which is predicted to be maximal at low glycolytic flux and high acetate concentration. Acetate should also increase the volumetric productivity (Figure 2G), which is maximal at high glycolytic flux and medium acetate concentration (20-30 mM), as acetate enhances both the production flux (Figure 2D) and the growth rate (Figure 2C).

These predictions suggest that acetate can enhance bioproduction through the combined effects of three distinct mechanisms: (i) by reducing acetate production through thermodynamic control of the Pta-AckA pathway, thereby minimizing carbon loss from the biosynthetic pathway, (ii) by replenishing the acetyl-CoA pool, which can increase the bioproduction flux through the acetyl-CoA-sensitive pathway, and (iii) by improving productivity by enhancing biomass accumulation. These mechanisms may work independently or synergistically to improve bioprocess efficiency.

### 3.3. Validation of model predictions for mevalonate production

We tested the predictions of the model for the production of mevalonate, a precursor for a broad range of high-value chemicals (e.g., carotenoids and steroids), pharmaceuticals (e.g., artemisinin), and biofuels (e.g., isoprenes) (C.-H. Wang et al., 2023). The mevalonate pathway is well understood, making it ideal for testing new strategies for bioprocess optimization that can readily be applied at the industrial scale (C.-H. Wang et al., 2023).

The first step of the mevalonate production pathway involves the conversion of acetyl-CoA into acetoacetyl-CoA by acetoacetyl-CoA thiolase (Figure 3A). The affinity of this enzyme for acetyl-CoA (K_M_ = 0.47 mM, Duncombe and Frerman, 1976) is low relative to the intracellular concentration of acetyl-CoA (0.02-0.6 mM). The production flux should therefore be responsive to the predicted increase in acetyl-CoA concentration caused by acetate supplementation (Figure 2E). Acetoacetyl-CoA is then converted into 3-hydroxy-3-methylglutaryl-CoA by HMG-CoA synthase, and finally into mevalonate by HMG-CoA reductase (Figure 3A). To test the predictions of the model against experimental data, we transformed *E. coli* K-12 BW25113 with the pMevT plasmid (Martin et al., 2003), which contains *atoB* (acetoacetyl-CoA thiolase), *HMGS* (HMG-CoA synthase), and *tHMGR* (truncated HMG-CoA reductase). We grew this strain on glucose, with or without acetate (30 mM), and measured growth rates, extracellular fluxes, and mevalonate bioproduction metrics.

**Figure 3.**
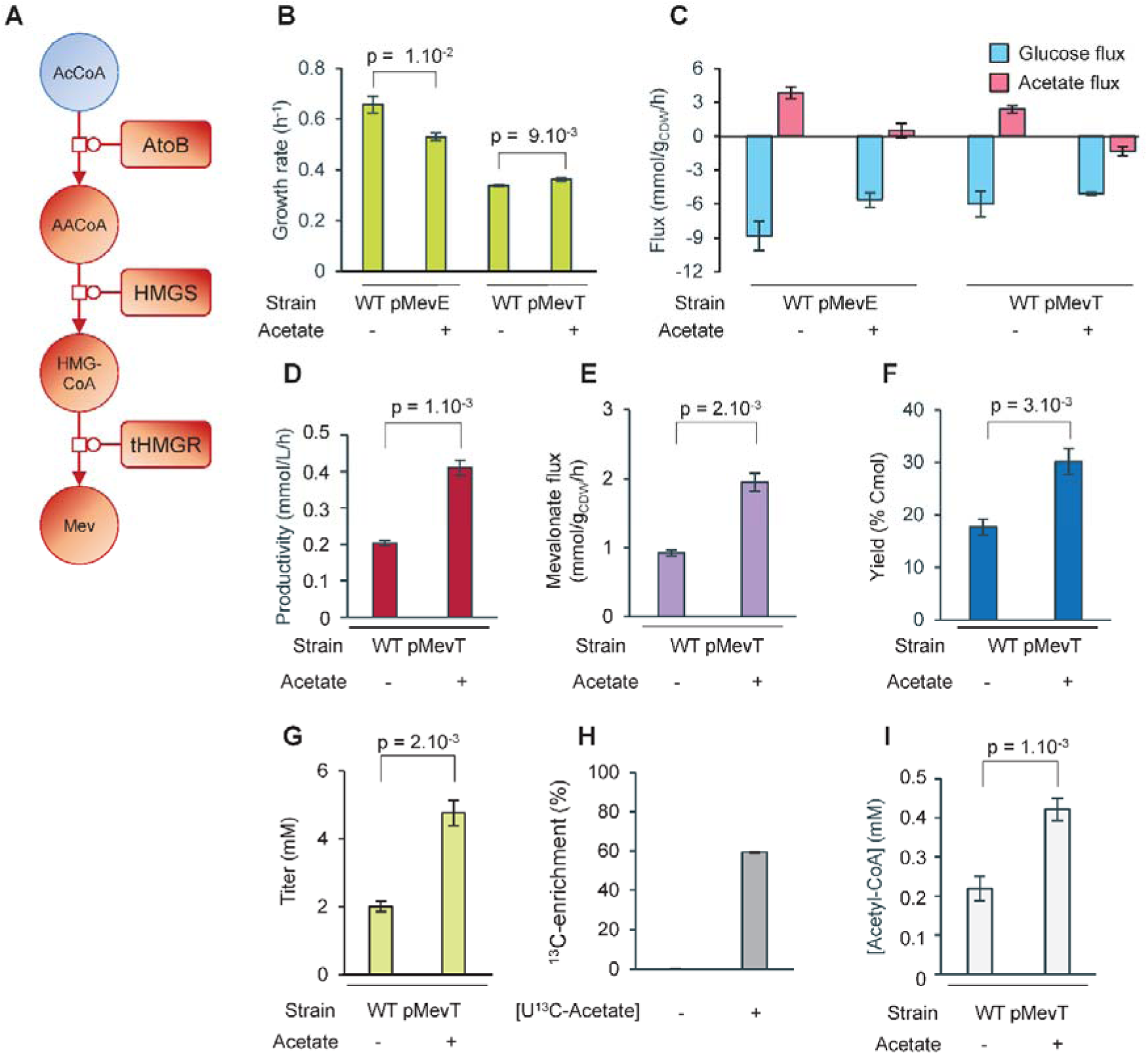
Impact of acetate on mevalonate production. (A) Representation of the synthetic mevalonate production pathway implemented in *E. coli* K-12 BW25113 pMevT. Growth rate (B), glucose and acetate fluxes (C), mevalonate production flux (D), yield (E), productivity (F), and titer (G) measured during growth of *E. coli* on glucose (15 mM) with or without 30 mM acetate. (H) ^13^C-enrichment on mevalonate produced on unlabeled glucose (15 mM) with or without 30 mM U-^13^C_2_-acetate. (I) Intracellular acetyl-CoA concentration measured in the producing strain during growth on glucose (15 mM) with or without 30 mM acetate. Mean values ± standard deviations (error bars) were estimated from three independent biological replicates.

In the absence of acetate, expression of the production pathway led to a 49% reduction in growth rate (from 0.66 to 0.33 h⁻¹, Figure 3B) while mevalonate was produced at 0.9 mmol/g_CDW_/h (Figure 3D). The reduced growth rate can be explained by the metabolic burden of heterologous protein expression (Snoeck et al., 2024), and by the high level of mevalonate production, which in this setting diverted 15% of carbon away from anabolism. The mevalonate-producing strain also produced acetate (2.4 mmol/g_CDW_/h; 13% of carbon uptake; Figure 3C), albeit at a significantly lower rate than the parent strain with an empty plasmid (3.8 mmol/g_CDW_/h).

In the presence of 30 mM acetate, the acetate flux was reversed (−1.3 mmol/g_CDW_/h, Figure 3C), leading to co-consumption of acetate with glucose, consistent with model predictions and results obtained in the non-producing parent strain under similar conditions (Millard et al., 2023). The glucose uptake rate decreased by 15% in the presence of acetate, in line with its known inhibitory effect on glucose uptake (Millard et al., 2021). Notably, mevalonate production increased significantly, with the production flux doubling from 0.9 to 2.0 mmol/g_CDW_/h (+112%, Figure 3D) and leading to a similar increase in volumetric productivity (+102%, Figure 3F), while the titer increased by 137% (Figure 3G). These results confirm that acetate enhances mevalonate production, as predicted by the model. Acetate supplementation also improved growth (Figure 3B), with the growth rate increasing slightly but significantly from 0.33 to 0.36 h⁻¹ (+8%), similar to previous observations (Millard et al., 2023). This result, also predicted by the model at low glycolytic flux, indicates that acetate partially alleviates the metabolic burden associated with the synthetic pathway and carbon diversion.

The mevalonate flux on glucose alone represents 47% of the flux on glucose and acetate, suggesting that acetate is responsible for 53% of the mevalonate flux. To confirm the contribution of acetate to mevalonate biosynthesis, we conducted a ^13^C-labeling experiment, growing the mevalonate-producing strain on unlabeled glucose and U-^13^C₂-acetate (Figure 3H). Under glucose-only conditions, the ^13^C enrichment of mevalonate was 0%, but in the presence of ^13^C-labeled acetate, enrichment increased to 59%, with only 41% of mevalonate derived from glucose. These values align closely with those estimated from the mevalonate flux measurements. Furthermore, the increase in mevalonate flux (from 0.9 to 2.0 mmol/g_CDW_/h, a difference of 6.6 Cmmol/g_CDW_/h) quantitatively matched the change in acetate balance (from +2.4 to –1.3 mmol/g_CDW_/h, a difference of 7.4 Cmmol/g_CDW_/h). These results demonstrate that the increase in mevalonate flux was driven exclusively by acetate.

We performed quantitative metabolomics experiments to verify the model prediction that the flux increase is caused by an increase in the intracellular concentration of acetyl-CoA. The acetyl-CoA concentration increased from 0.22 mM in the absence of acetate to 0.42 mM in the presence of 30 mM acetate (Figure 3I), in agreement with model predictions. Given the *in vitro* K_M_ of acetoacetyl-CoA thiolase for acetyl-CoA (0.47 mM), this concentration increase should theoretically translate into a 68% increase in flux. While qualitatively consistent with experimental flux measurements, the actual flux increase (+112%) was much higher than this metabolomics-derived estimate. This suggests that the *in vivo* K_M_ of acetoacetyl-CoA thiolase is higher than measured *in vitro*, as observed for other enzymes in a recent *in vivo* enzymology study (Castaño-Cerezo et al., 2024). Acetate may also enhance expression and thus activity in the heterologous pathway, as reported previously for the expression of the growth machinery (Millard et al., 2023).

These results highlight the value of acetate for bioproduction with an acetyl-CoA-sensitive pathway, as simply adding 30 mM acetate increased the mevalonate production flux by 112%, productivity by 102%, and titer by 137%. The findings indicate that acetate enhances mevalonate production purely at the metabolic level, primarily through an acetate-induced increase in the acetyl-CoA pool. No acetate is produced, and acetate is almost entirely converted into mevalonate, with the beneficial side-effect of a slight but significant increase in growth.

### 3.4. Predictions for an acetyl-CoA-insensitive production pathway

Following the same strategy, we investigated an alternative scenario in which the production pathway operates under saturation (K_M_ = 5 µM) and is insensitive to changes in acetyl-CoA induced by acetate. In this case, the model predicts that the growth physiology of *E. coli* (glucose uptake flux, acetate flux and growth rate, Figure 4A-C) should respond to perturbations similarly as in the non-producing strain and as in the producing strain with an acetyl-CoA-sensitive pathway, albeit with different trends for the three process efficiency metrics. Productivity (Figure 4C) is predicted to be maximal at medium acetate concentrations (5–10 mM), where acetate stimulates growth. The production flux (0.8–1 mmol/g_CDW_/h, Figure 4D) remains close to the maximal enzyme capacity (V_max_ = 1 mmol/g_CDW_/h), confirming optimal enzyme utilization, though the flux is less responsive to changes in acetyl-CoA concentrations (Figure 4E), as expected. Interestingly, the yield is highest at the lowest glucose uptake rate (Figure 4F), a trend commonly reported in production strains (Kyselova et al., 2018; Lim and Jung, 2017; Zhuang et al., 2013). Acetate is predicted to enhance productivity at all glycolytic activity levels (Figure 4G) but does not improve the product yield as strongly as it does in the acetyl-CoA sensitive pathway.

**Figure 4.**
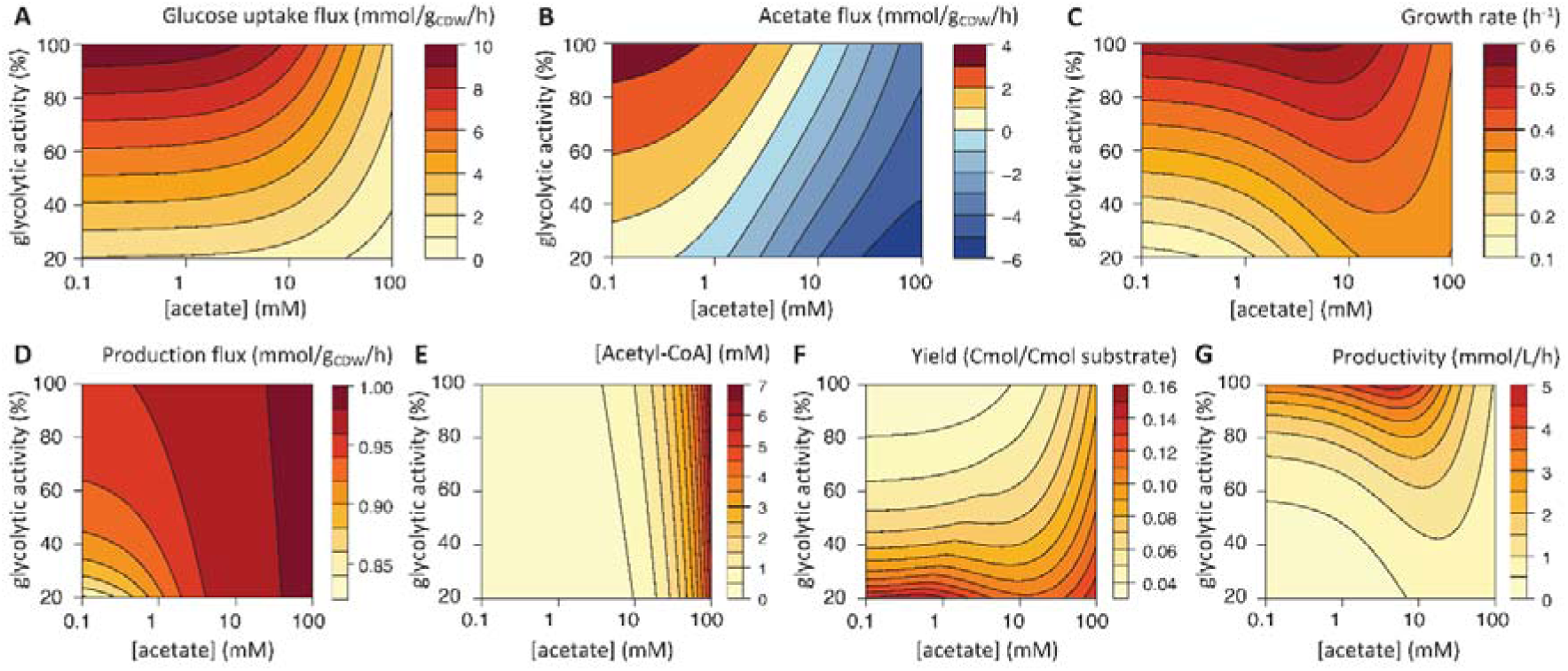
Predicted response to glycolytic and acetate perturbations of an *Escherichia coli* strain with an acetyl-CoA-insensitive production pathway. Glucose uptake flux (A), acetate flux (B), growth rate (C), product flux (D), intracellular acetyl-CoA concentration (E), product yield (F) and volumetric productivity (G) simulated over a broad range of glycolytic activity levels (20 to 100% of the initial V_max_) and acetate concentrations (0.1 to 100 mM).

These predictions suggest that even when using an acetyl-CoA-insensitive production pathway, acetate can still enhance bioproduction. While the two key benefits observed in acetyl-CoA-sensitive pathways – related to the reduction of acetate overflow and increase in intracellular acetyl-CoA concentration – are no longer present, acetate can still boost productivity by promoting biomass accumulation.

### 3.5. Validation of model predictions for 3-hydroxypropionate production

We validated this model by comparing predicted and measured production parameters for 3HP, an important platform chemical in biotechnology with various applications in bioplastics (e.g., poly-3-hydroxypropionate), chemicals (e.g., acrylic acid and 1,3-propanediol), and pharmaceuticals (e.g., beta-lactones) (Vidra and Németh, 2017). The first step of the 3HP production pathway involves the conversion of acetyl-CoA into malonyl-CoA by a native acetyl-CoA carboxylase (Figure 5A). This enzyme has a high affinity for acetyl-CoA (K_M_ = 18 µM, (Soriano et al., 2006)) and is therefore expected to operate near saturation under physiological acetyl-CoA concentrations. Malonyl-CoA is then converted into 3HP by malonyl-CoA reductase (Figure 5A).

**Figure 5.**
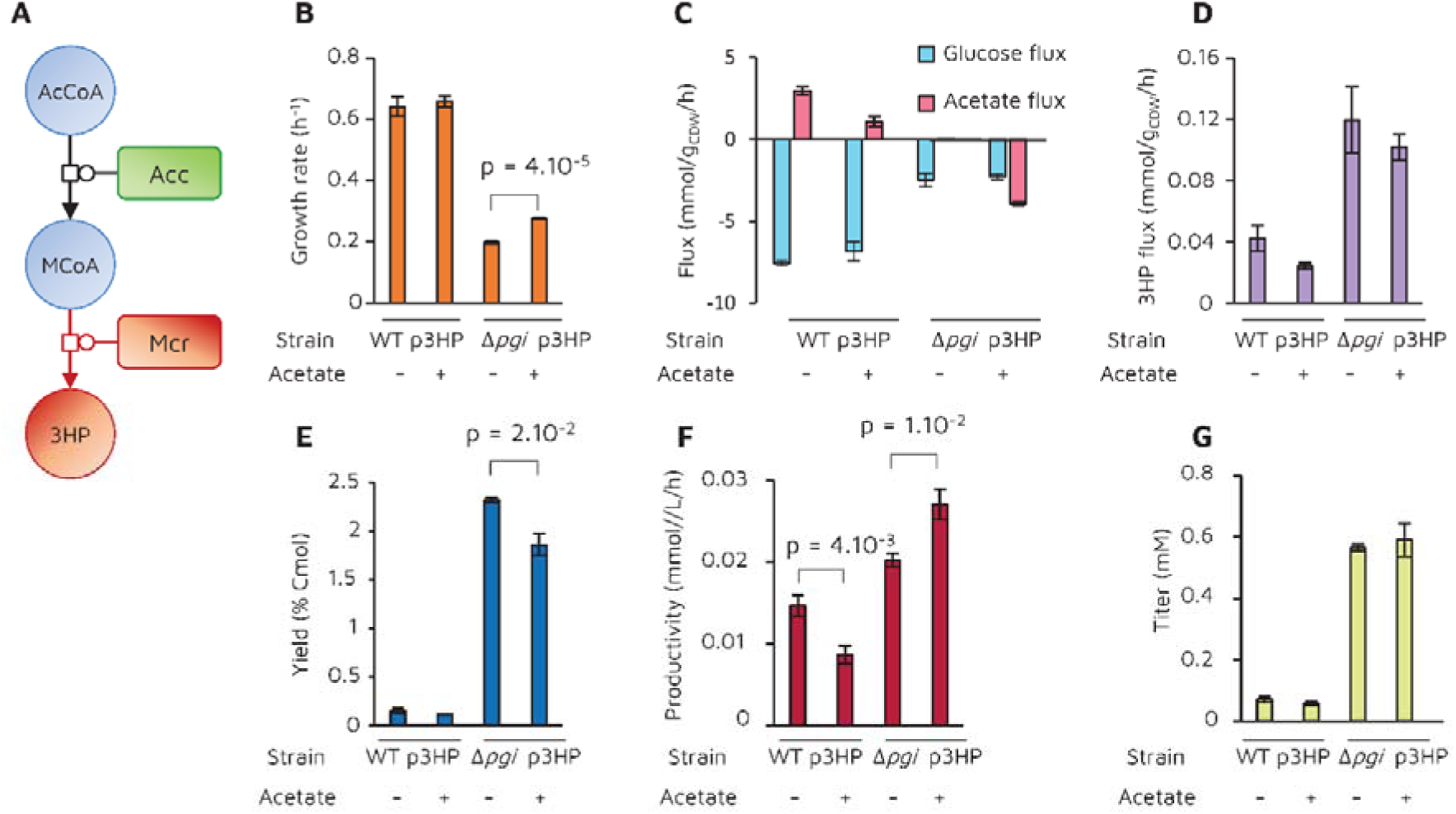
Impact of acetate on 3-hydroxypropionate production in parent (WT) and Δ*pgi E. coli* strains. (A) Representation of the 3-hydroxypropionate production pathway implemented in *E. coli*. The native metabolism is shown in blue (metabolites) and green (enzymes), and the heterologous reaction is shown in red. (B) Growth rate, (C) glucose and acetate fluxes, (D) 3-hydroxypropionate production flux, (E) yield, (F) volumetric productivity and (G) titer measured during growth of *E. coli* on glucose (15 mM) with or without 10 mM acetate. Mean values ± standard deviations (error bars) were estimated from three independent biological replicates.

We performed growth experiments on glucose, with or without acetate, with an *E. coli* K-12 BW25113 strain transformed with p3HP, a plasmid carrying *mcr* (malonyl-CoA reductase). Since the model predicted that optimal productivity would be achieved at slightly lower acetate concentrations than for acetyl-CoA-sensitive production pathways, we used an acetate concentration of 10 mM. Since malonyl-CoA reductase requires two NADPH molecules, we also performed production experiments with a Δ*pgi* strain, which is expected to increase NADPH availability, thereby enhancing flux through NADPH-dependent biosynthetic pathways (Chemler et al., 2010).

In the absence of acetate, the growth of the modified strain did not differ from that of the parent strain (Figure 5B-C, with control experiments shown in Supplementary Figure S1). Deletion of *pgi* reduced growth and glucose uptake rates (−69 and −67%, respectively) and abolished acetate production (Figure 5B-C), consistent with previous reports (Long and Antoniewicz, 2019; Millard et al., 2023). Despite its negative impact on *E. coli* physiology, deletion of *pgi* led to a 183% increase in 3HP production flux (Figure 5D), a 16-fold increase in yield (Figure 5E), a 38% increase in volumetric productivity (Figure 5F), and an 8-fold increase in titer (Figure 5G), confirming that the Δ*pgi* strain is advantageous for the production of NADPH-dependent compounds.

In the 3HP-producing parent strain, acetate supplementation led to reduced acetate production (−64%, Figure 5C), in agreement with model predictions (Figure 4B). However, 3HP production decreased, in terms of production flux by 42%, yield by 26%, productivity by 41%, and titer by 20%. These results are consistent with previously reported negative effects of acetate on bioproduction (Eiteman and Altman, 2006). Interestingly, the model had predicted a slight increase in yield and productivity under high glycolytic flux; this discrepancy indicates that the predictive accuracy of the model for these parameters is lower in this specific context. In the Δ*pgi* strain, acetate supplementation led to co-utilization of acetate with glucose, in line with model predictions (Figure 4B). While acetate had no effect on the 3HP production flux, it significantly enhanced growth (+40%), consistent with predictions for a production pathway operating at saturation and under low glycolytic flux. The production yield was reduced (−20%, Figure 5E), as the additional carbon from acetate was used exclusively for anabolic processes, as also predicted by the model (Figure 4F). However, the faster biomass accumulation resulted in a substantial increase in productivity (+34%, Figure 5F), again consistent with model predictions (Figure 4G). Therefore, in the presence of acetate, the productivity of the Δ*pgi* strain was higher than that of the producing parent strain, despite a pronounced reduction in growth rate.

These results demonstrate that by promoting biomass accumulation and overall productivity, acetate supplementation also enhances the bioproduction of acetyl-CoA-derived compounds when the pathway is operating near saturation.

## 4. Discussion

Despite recent discoveries suggesting that acetate may benefit *E. coli*, acetate is still widely regarded as a problem in bioprocesses. In this study, we used kinetic modeling to explore the effects of acetate in glucose-based bioproduction. Our results indicate that while acetate is indeed detrimental to bioprocesses when glycolytic flux is high, it can be beneficial at low glycolytic flux. Low glycolytic flux conditions are common in industrial bioprocesses because of the intrinsically lower glucose uptake rates of engineered strains or due to cultivation strategies that reduce the glycolytic flux (e.g., fed-batch and chemostat cultures) (Carneiro et al., 2013). These findings suggest that acetate could serve as a metabolic booster for many *E. coli*-based bioprocesses, optimizing the fundamental trade-off between growth and production, a major constraint on bioprocess performance (Mannan et al., 2025).

Simulations indicate that acetate’s positive impact is mediated by three distinct mechanisms: (i) the presence of acetate minimizes carbon loss from the biosynthetic pathway by reducing acetate production, (ii) acetate enhances the bioproduction flux of acetyl-CoA-derived compounds by increasing the acetyl-CoA concentration, and (iii) acetate improves volumetric productivity by promoting biomass accumulation. Experimental results confirm these mechanisms and demonstrate that they collectively enhance bioprocess efficiency. These findings are directly transferable to industrial bioprocesses based on *E. coli*.

Results also show that the relative contribution of each mechanism to bioprocess improvement varies depending on the strain’s genetic background and the synthetic pathway involved. For mevalonate production, an acetyl-CoA sensitive pathway, all three mechanisms contributed, with the main benefit being the increase in biosynthetic flux, driven by reduced acetate accumulation and an increased acetyl-CoA pool. The growth-promoting effect of acetate played only a minor role. In contrast, in 3HP production, an acetyl-CoA insensitive pathway, no flux increase was observed in either the parent or Δ*pgi* producing strains, and the increase in productivity observed in the Δ*pgi* strain was entirely due to acetate’s positive effect on growth.

Understanding how enzyme saturation shapes microbial production fluxes and growth provides insights to optimize bioproduction processes. Our findings demonstrate that acetate can help balance cell growth and production, a common challenge in engineered strains. The beneficial effect of acetate, validated here on glucose, may also extend to other industrially relevant glycolytic substrates, such as glycerol (Millard et al., 2023). Furthermore, since the positive effect of acetate on growth under conditions of low glycolytic flux appears to be general, productivity improvements may be achievable for metabolic products not derived from acetyl-CoA—an area that warrants further investigation.

The modeling approach developed in this study is a valuable tool for the rational optimization of *E. coli*-based bioprocesses and can be extended to assess the role of acetate in other metabolic contexts. Our findings demonstrate how acetate, traditionally seen as Mr. Hyde, sabotaging productivity, can become Dr. Jekyll, enhancing bioprocess efficiency. We hope this work helps highlight the dual nature of acetate, which is key to understanding its role and unlocking its full potential in biotechnology. Since acetate can be sourced from CO_2_ and various waste streams, using it as a metabolic booster in bioproduction processes is also a means to improve their environmental sustainability.

## CRediT authorship contribution statement

**Thomas Gosselin-Monplaisir:** Conceptualization, Methodology, Investigation, Software, Formal analysis, Visualization, Writing – review & editing. **Denis Jallet:** Methodology, Investigation, Writing – review & editing. **Erwana Harscoet:** Methodology, Investigation, Writing – review & editing. **Sandrine Uttenweiler-Joseph:** Conceptualization, Writing – review & editing. **Stéphanie Heux:** Conceptualization, Funding acquisition, Supervision, Methodology, Writing – review & editing. **Pierre Millard:** Conceptualization, Funding acquisition, Project administration, Supervision, Methodology, Software, Formal analysis, Visualization, Writing – original draft, Writing – review & editing.

## Declaration of competing interest

The authors declare no conflict of interest.

## Funding

The PhD thesis of Thomas Gosselin-Monplaisir was funded by the Region Occitanie and the MICA department of INRAE (grant COCA-COLI). Thomas Gosselin-Monplaisir also benefited from a grant managed by the Agence Nationale de la Recherche under the Investissements d’Avenir Programme (grant ANR-18-EURE-0021).

## Acknowledgements

The authors thank MetaboHub-MetaToul (Metabolomics & Fluxomics facilities, Toulouse, France, https://mth-metatoul.com), which is part of the French National Infrastructure for Metabolomics and Fluxomics (https://www.metabohub.fr), funded by the ANR (MetaboHUB-ANR-11-INBS-0010), for access to MS and NMR facilities. The authors also thank Hanna Kulyk, Maud Heuillet, and Nina Lager-Lachaud (MetaboHub-MetaToul, Toulouse Biotechnology Institute, Toulouse, France) for their help with MS analyses and data processing, and Sara Castaño Cerezo (Toulouse Biotechnology Institute, Toulouse, France) for insightful discussions.

## Notes

https://github.com/MetaSys-LISBP/glucose_acetate_bioproduction

### Competing Interest Statement

The authors have declared no competing interest.

