## Supplementary Figure 1 for "Acetate as a metabolic booster for glucose-based bioproduction in Escherichia coli"

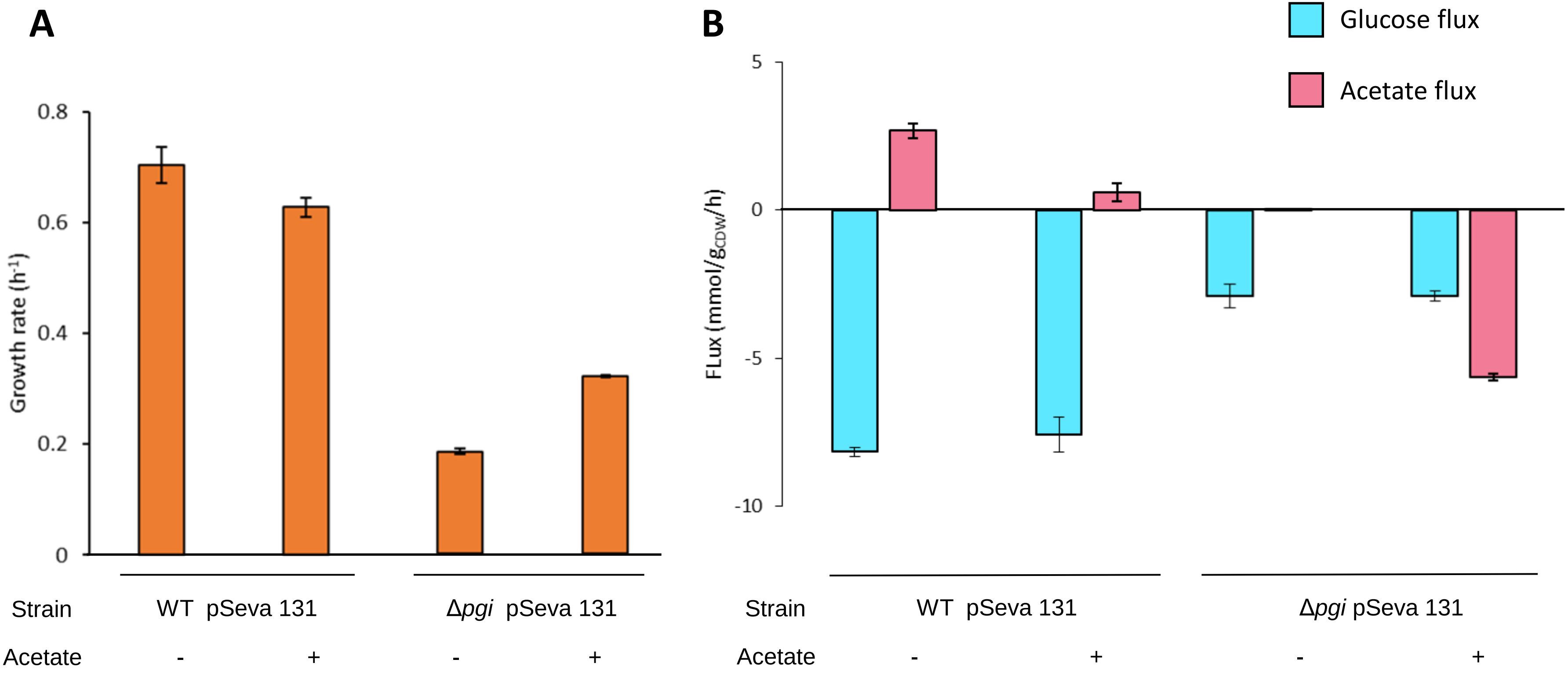


**Supplementary Figure 1.** Impact of acetate on *E. coli* WT and Δ*pgi* strains transformed with pSEVA 131: (A) growth rate, (B) glucose and acetate fluxes.
